# Intracellular stiffening competes with cortical softening in glioblastoma cells

**DOI:** 10.1101/2020.09.02.279794

**Authors:** Charlotte Alibert, David Pereira, Nathan Lardier, Sandrine Etienne-Manneville, Bruno Goud, Atef Asnacios, Jean-Baptiste Manneville

**Author notes:** these authors contributed equally (Charlotte Alibert, David Pereira). Author contributions: CA, DP, and NL performed experiments; SEM, BG, AA and JBM designed the project and conceived the experiments; CA, DP and JBM wrote the paper.

## Abstract

Cancer cell softening increases with the progression of the disease, suggesting that mechanical phenotyping could be used as a diagnostic or prognostic method. Here we investigate the cell mechanics of gliomas, brain tumors that originate from glial cells or glial progenitors. We use two microrheology techniques, a single cell parallel plates rheometer to probe whole cell mechanics and optical-tweezers to probe intracellular rheology. We show that cell mechanics discriminates human glioma cells of different grades. When probed globally, grade IV glioblastoma cells are softer than grade III astrocytoma cells, while they are surprisingly stiffer at the intracellular level. We explain this difference between global and local intracellular behaviours by changes in the composition and spatial organization of the cytoskeleton and by changes in nuclear mechanics. Our study highlights the need to combine rheology techniques for potential diagnostic or prognostic methods based on cancer cell mechanophenotyping.

## Introduction

Solid tumors are often first detected by palpation, indicating that they are more rigid than the surrounding tissues. In contrast, cancer cells appear softer than normal cells in many cancer types such as breast, kidney, prostate, or bladder cancers (Abidine et al., 2015; Alibert et al., 2017; Faria et al., 2008; Lekka et al., 1999; Rebelo et al., 2013; Thoumine and Ott, 1997; Ward et al., 1991). In addition, cancer cell softening has been reported to increase as the disease progresses and metastasizes (Alibert et al., 2017; Guck et al., 2005; Holenstein et al., 2019; Mandal et al., 2019), suggesting that cell mechanics could be used as a new diagnosis and/or prognosis tool, complementary to more classical molecular markers-based techniques.

Cancer cell softening may facilitate cell migration when cancer cells invade the confined space surrounding the tumor and move over long distances to form metastases. Why and how cells soften during tumor progression is still not clear and may result from a combination of numerous factors, among which modifications in the cell internal architecture, the cytoskeleton, the mechanical properties of the plasma membrane and its associated cortex or of the nucleus, cell contractility, signaling pathways and metabolism (Alibert et al., 2017). For instance, nuclear mechanics was recently shown to regulate the ability of cells to migrate through narrow pores or constrictions (Denais et al., 2016; McGregor et al., 2016; Raab et al., 2016) and to correlate with the invasiveness of cancer cells (Darling and Di Carlo, 2015; Denais and Lammerding, 2014; Gal and Weihs, 2012; Ketene et al., 2012; Xu et al., 2012).

We focus here on the mechanics of glioma cells. Gliomas are brain tumors originating from glial cells, principally oligodendrocytes and astrocytes (Maher et al., 2001), from glial progenitors and neural stem cells (Lee et al., 2018). Gliomas are classified in four grades (Louis et al., 2016, 2007) with grade III astrocytomas and grade IV glioblastomas being the most common and aggressive form of gliomas in adults and children. Despite the identification of several histological signatures, molecular markers such as EGFR, Sox2, PDGFRA, Olig2 and the two intermediate filament proteins GFAP and nestin (Ehrmann et al., 2005; Jacque et al., 1978; Quick et al., 2015; Tampaki et al., 2014; Yang et al., 2008), mutations such as IDH1/2 or perturbed signaling pathways such as the RTK/RAS/PI3K, the TP53 or RB pathways (Louis et al., 2016; Patel et al., 2014; Verhaak et al., 2010; Xie et al., 2015), the prognosis of gliomas remains poor. Even after treatment with combined chemotherapy, radiotherapy and surgery, the prognosis of patients with grade IV glioblastomas is low and survival rarely exceeds 15 months after diagnosis (Lieberman, 2017). Developing new techniques based on cell mechanics could thus be of especially great value in the case of gliomas.

Brain is the softest tissue in the organism with typical values of the Young modulus around 100 Pa up to a few kPa depending on the technique used (Budday et al., 2015; Lu et al., 2006; Reiss-Zimmermann et al., 2015). Consistently, cells from the central nervous system, glial cells and neurons, are more deformable than most eukaryotic cells, with glial cells being even softer than neurons (Lu et al., 2006), suggesting that glial cells provide a protective and compliant environment for neurons. In contrast with most solid tumors, gliomas appear to be softer than the surrounding healthy environment in which they develop, as shown *in vivo* using Magnetic Resonance Elastometry (MRE) (Reiss-Zimmermann et al., 2015; Streitberger et al., 2014). Atomic Force Microscopy (AFM) experiments showed that glioblastoma cells were softer than normal astrocytes (Pogoda et al., 2014).

Here we investigate the mechanical properties of glioma cells of different grades by probing whole cell rheology using a single cell uniaxial rheometer and intracellular rheology using optical tweezers combined with adhesive micropatterns. We show that, while glioma cells are softer than primary astrocytes at both whole cell and intracellular scales, human grade IV cells are softer than grade III cells only at the whole cell scale. Indeed, at the intracellular scale, grade IV cells are surprisingly stiffer than grade III cells. We rationalize these findings by showing differences in the spatial organization of the cytoskeleton in grade III and grade IV glioma cells, and by identifying nuclear mechanics as a major contributor to intracellular stiffening in grade IV glioma cells as compared to grade III cells.

## Results

### Rat glioblastoma cells are softer than rat primary astrocytes

To test whether glioma cells are softer than their normal counterparts, we first compared the mechanics of rat glioblastoma F98 cells (grade IV) with that of primary rat astrocytes Using the uniaxial single cell microplate rheometer, in which single cells are subjected to oscillatory uniaxial compressions between two parallel glass microplates (Fig. S1A), we show a two-fold decrease in the complex shear modulus in glioblastoma cells as compared to astrocytes (Fig. 1A,B), with a decrease both in the elastic storage modulus *G*′ and in the viscous loss modulus *G*″ (Fig. 1B). Because the amplitude of the oscillations is about 1 μm, cytoplasmic organelles are not significantly compressed and the cell cortex is certainly the main contributor to the mechanical moduli measured in this experiment as previously discussed (Durand-Smet et al., 2014; Wu et al., 2018). Our results thus show that the cortical stiffness of glioma cells is decreased compared to that of normal glial cells. Note that in all cases the storage modulus dominates the loss modulus, indicating that elasticity contributes more than viscosity to the complex shear modulus, as we and others reported previously for other cell types (Bufi et al., 2015b; Lu et al., 2006; Mandal et al., 2019, 2016).

**Figure 1.**
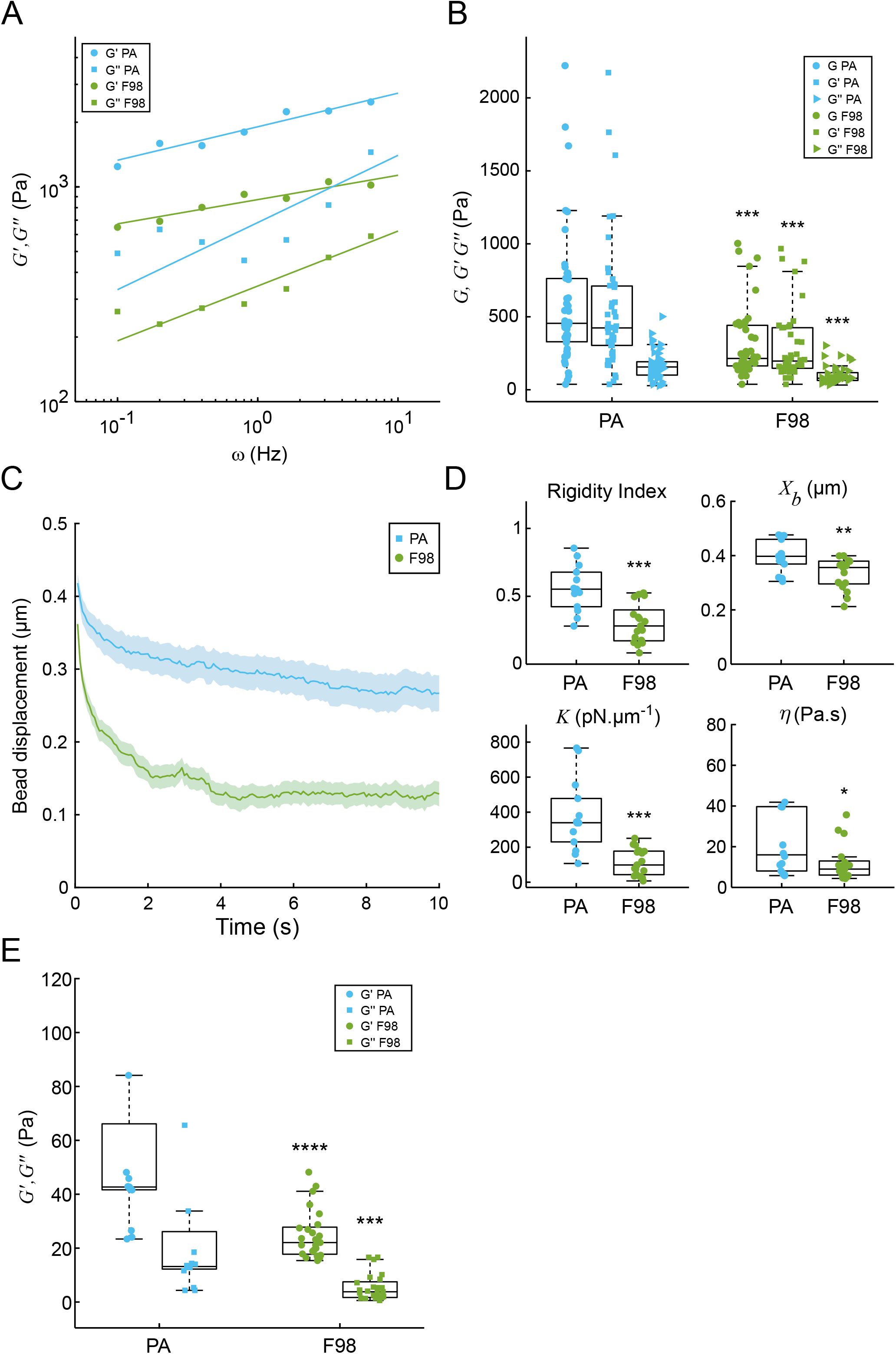
Rat glioblastoma cells are softer than rat primary astrocytes at the whole cell scale and at the intracellular scale. A-B. Whole cell microrheology with a uniaxial rheometer. A. Storage *G*′ (circles) and loss *G*″ (squares) moduli as a function of the strain frequency *ω* of a single primary rat astrocyte (‘PA’, blue) and rat F98 glioma cell (‘F98’, green). Both moduli were fitted using a power law (dashed line). B. Averaged values of the amplitude of the complex shear modulus (*G*_0_), the storage (*G*′) and the loss (*G*″) moduli of primary rat astrocytes and of rat glioblastomas F98 cells at the scale of the whole cell with the single cell microplate rheometer. The shear modulus gives an effective cellular rigidity, while storage and loss modulus respectively quantify elastic and viscous cell behaviors. Data were obtained from n=47 astrocytes and n= 42 F98 cells. C-E. Intracellular microrheology by optical tweezers-based viscoelastic relaxation experiments. C. Averaged relaxation curves of the position of an internalized bead towards the trap center following a 0.5 μm step displacement of the cell. Relaxations in rat primary astrocytes and in rat glioblastomas F98 cells are shown in blue and in green respectively. Data were obtained from n=17 and 25 beads in astrocytes and in F98 cells, respectively. D. Quantification of the relaxation curves shown in C using the phenomenological and the viscoelastic methods. The bead step displacement *X*_*b*_ and the rigidity index *RI* are two phenomenological parameters corresponding to the initial bead displacement in the optical trap and to the normalized area below the relaxation curve respectively. The spring constant *K* and the viscosity *η* are obtained by fitting the relaxation curves with the standard linear liquid (SLL) model. Data were obtained from n=14 and 17 beads in astrocytes and in F98 cells, respectively. E. Quantification of the relaxation curves shown in C using the power-law method. The components of the complex shear modulus, *G*′ and *G*″, are obtained by fitting the relaxation curves with power-law model as in (Mandal et al., 2016). Data were obtained from n=16 and 25 beads in astrocytes and in F98 cells, respectively.

We then used an active microrheology technique based on the coupling of optical tweezers with fast confocal microscopy (Fig. S1B) (Guet et al., 2014; Mandal et al., 2016) to measure intracellular viscoelastic properties. Beads (2 μm diameter) internalized by the cells are trapped by the optical tweezers. The cell is then displaced in a stepwise fashion and the relaxation of the bead towards the trap center is recorded. The viscoelastic relaxation curves are analyzed by three methods (see Materials and Methods). First, a phenomenological model-independent approach yields two parameters, the initial bead step displacement *X*_*b*_ and the rigidity index *RI* which both increase when the rigidity of the bead microenvironment increases. Second, a fit using the standard linear liquid (SLL) viscoelastic model yields the spring constant *K* and the dynamic viscosity *η*. Third, assuming that the cell cytoplasm obeys power-law rheology, fitting the curves as in (Mandal et al., 2019, 2016) gives the storage modulus *G*′ and the loss modulus *G*″. Our results show that rat glioblastoma F98 cells are softer than normal rat astrocytes at the intracellular level. The bead relaxes much faster and closer to the trap center in glioblastoma cells (Fig. 1C). The values of all the parameters extracted from the relaxation curves decrease significantly indicating a lower rigidity in glioblastoma cells originating both from a decrease in elasticity and in viscosity (Fig. 1D,E).

### Grade IV human glioma cells are softer than grade III cells at the whole cell scale but stiffer at the intracellular scale

We next compared the mechanics of two human glioma cell lines of different grades, the grade III astrocytoma U373 cell line (Westermark et al., 1973) and the grade IV glioblastoma U87 cell line (Allen et al., 2016). Grade III and grade IV glioma cell lines were shown previously to exhibit different migratory properties (Camand et al., 2012; Sabari et al., 2011), suggesting that they may have different mechanical properties. Using the single cell microplate rheometer (Fig. S1A and S2), we found that grade IV cells are softer than grade III cells. The frequency-dependent shear modulus *G*_0_ (*ω*) was lower in grade IV cells (Fig. 2A), with a 2.5-fold reduction of the elastic storage modulus *G*’ and a 1.8-fold reduction in the viscous loss modulus *G*’’ (Fig. 2B).

**Figure 2.**
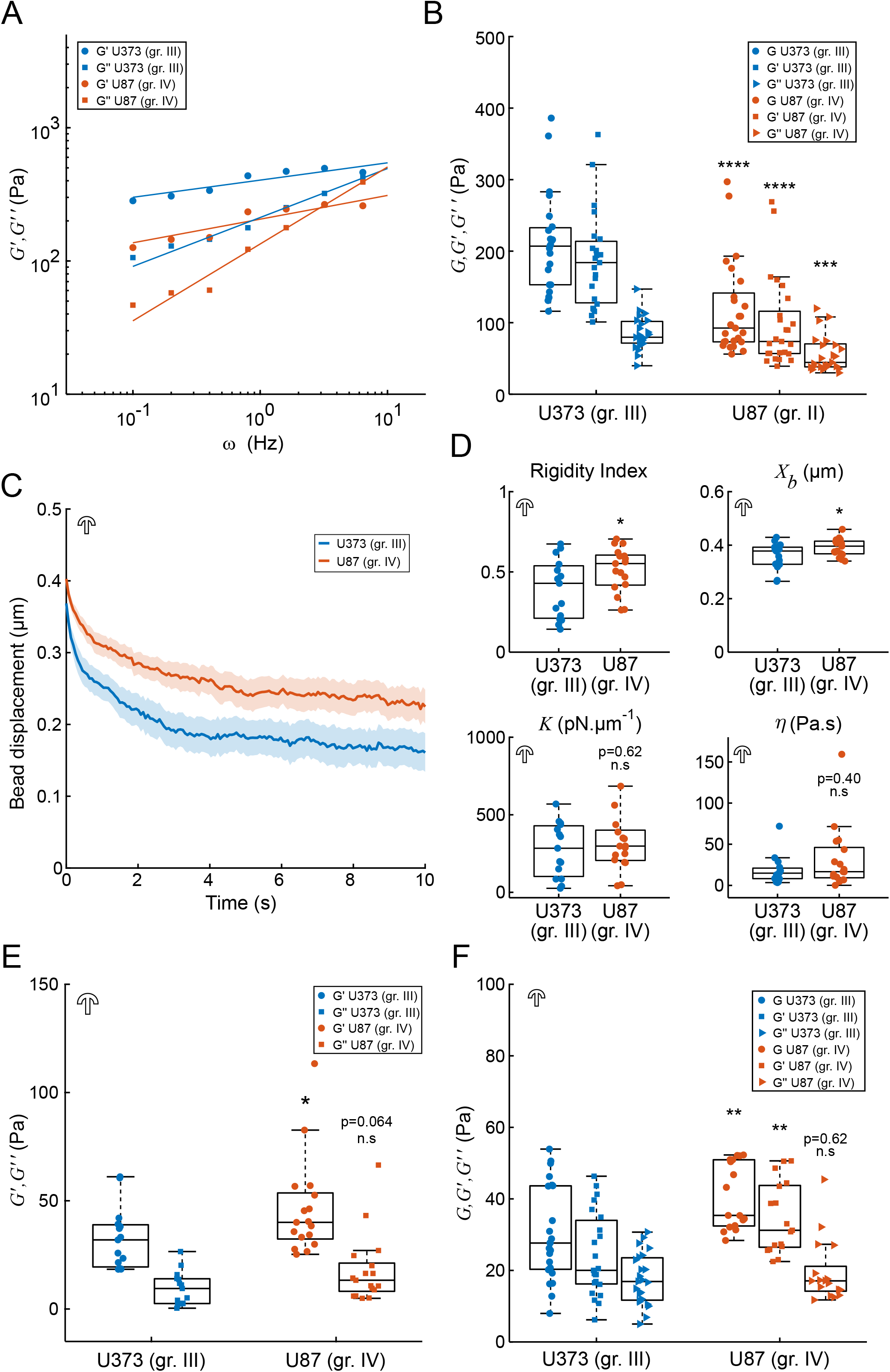
Grade IV human glioma cells are softer than grade III cells at the cortical scale but stiffer at the intracellular scale. A-B. Whole cell microrheology with a uniaxial rheometer. A. Storage (*G*′, circles) and loss (*G*″, squares) moduli as a function of the strain frequency *ω* of a typical grade III and grade IV cells (blue and orange respectively). The grey line is a power-law fit of the data. B. Averaged values of the amplitude of the complex shear modulus (*G*_0_), the storage (*G*′) and the loss (*G*″) moduli of grade III and grade IV glioma cells. Data were obtained from n=23 and 25 grade III and grade IV cells, respectively. C-F. Intracellular microrheology by optical tweezers-based viscoelastic relaxation and oscillatory experiments. C. Averaged relaxation curves for grade III glioma cells (blue) and for grade IV glioma cells (orange). Cells were plated on crossbow-shaped micropatterns adapted to each cell type (see Materials and Methods and Fig. S3). Data were obtained from n=22 and 17 grade III and grade IV cells, respectively. D. Phenomenological parameters (bead step amplitude *X*_*b*_ and rigidity index *RI*) and parameters obtained by fitting the relaxation curves with the SLL model (spring constant *K* and viscosity *η*). Data were obtained from n=15 and 17 grade III and grade IV cells, respectively. E. Fit parameters from the power-law analysis (elastic storage modulus *G*′, viscous loss modulus *G*″). Data were obtained from n=15 and 17 grade III and grade IV cells, respectively. F. Amplitude of the complex shear modulus and storage *G*′ and loss moduli *G*″ measured by oscillatory active microrheology in grade III and grade IV glioma cells plated on crossbow-shaped micropatterns. Data were obtained from n=23 and 17 grade III and grade IV cells, respectively.

We then asked whether the cortical softening of grade IV cells observed at the scale of the whole cell was paralleled by an intracellular softening. Intracellular relaxation experiments showed no significant difference when comparing freely adhering human grade III and grade IV glioma cells (Fig. S3) with either relaxation rheology (Fig. S1B, Fig. S3A-C) or oscillatory rheology (Fig. S1C, Fig. S3D). However, in these conditions, cells exhibit variable morphologies and adhesive properties. To obtain more robust measurements, we immobilized cells on crossbow-shaped adhesive micropatterns, which induces a reproducible spatial organization of the cytoskeleton and of intracellular organelles (Schauer et al., 2010; Théry et al., 2006) and allows spatial mapping of the mechanical properties of the cell interior (Mandal et al., 2016). We designed adhesive fibronectin-coated crossbow-shaped micropatterns adapted to grade III U373 cells and to grade IV U87 cells (Fig. S4). Measurements of the cell spreading area on fibronectin (Fig. S4A,B) and of the cell volume (Fig. S4C,D) showed that U373 cells have a 1.66-times larger volume *V* and a 1.5-larger spreading area *S* than U87 cells. The averaged aspect ratio α = *V*^1/3^/*S*^1/2^, which indicates if the cells display a globular (α ≳ 1) or a flattened (α < 1) shape, was the same for both cell types (α = 0.37), indicating that the ratio between cell height and the typical length scale of the cell-substrate contact area is similar for both U87 and U373 cell types. Thus, we used micropatterns of area equal to the mean spreading area of freely adhering cells (Fig. S4E,F). Relaxation and oscillatory active rheology experiments showed that, surprisingly, U87 grade IV cells are stiffer than U373 grade III cells at the intracellular scale (Fig. 2C-F). Relaxation was slower in grade IV cells (Fig. 2C) and quantification showed an increase in the phenomenological parameters (initial bead step amplitude *X_b_* and rigidity index *RI*) (Fig. 2D), and in the shear modulus deduced from a power-law analysis (Fig. 2E). We also measured a small but not statistically significant increase in the spring constant and in the viscosity (*K* and *η*) deduced from a fit of the relaxation curves using the SLL model. Similar results were obtained with intracellular oscillatory rheology (Fig. 2F, Fig. S5) with a significant increase in the elastic storage modulus *G*′, confirming the increased intracellular rigidity in grade IV cells. We conclude that grade IV glioma cells are softer than grade III glioma cells at the whole cell scale (Fig. 2A,B), in agreement with data published in other cancer types (Alibert et al., 2017), while their intracellular rigidity is significantly higher (Figs. 2C-F).

### The expression levels of cytoskeletal proteins are lower in grade IV glioblastoma cells

The cytoskeleton, including actin microfilaments, microtubules and intermediate filaments, plays a critical role in both cortical and intracellular mechanics. To rationalize the differences in glioma cell mechanics, we have first quantified the levels of cytoskeletal proteins in both cell types using Western blot. We found that the total levels of actin and tubulin, and of the three intermediate filament proteins vimentin, GFAP and nestin, are reduced to different extent in grade IV cells as compared to grade III cells when measured by Western blot (Fig. 3). The decrease in actin or vimentin levels was not statistically significant. In contrast, the expression levels of tubulin, GFAP and nestin were strongly reduced in grade IV cells. These results show that the amount of cytoskeletal proteins is lower in grade IV cells, consistent with the whole cell softening of grade IV cells measured with the uniaxial parallel-plates single cell rheometer (Fig. 2A,B).

**Figure 3.**
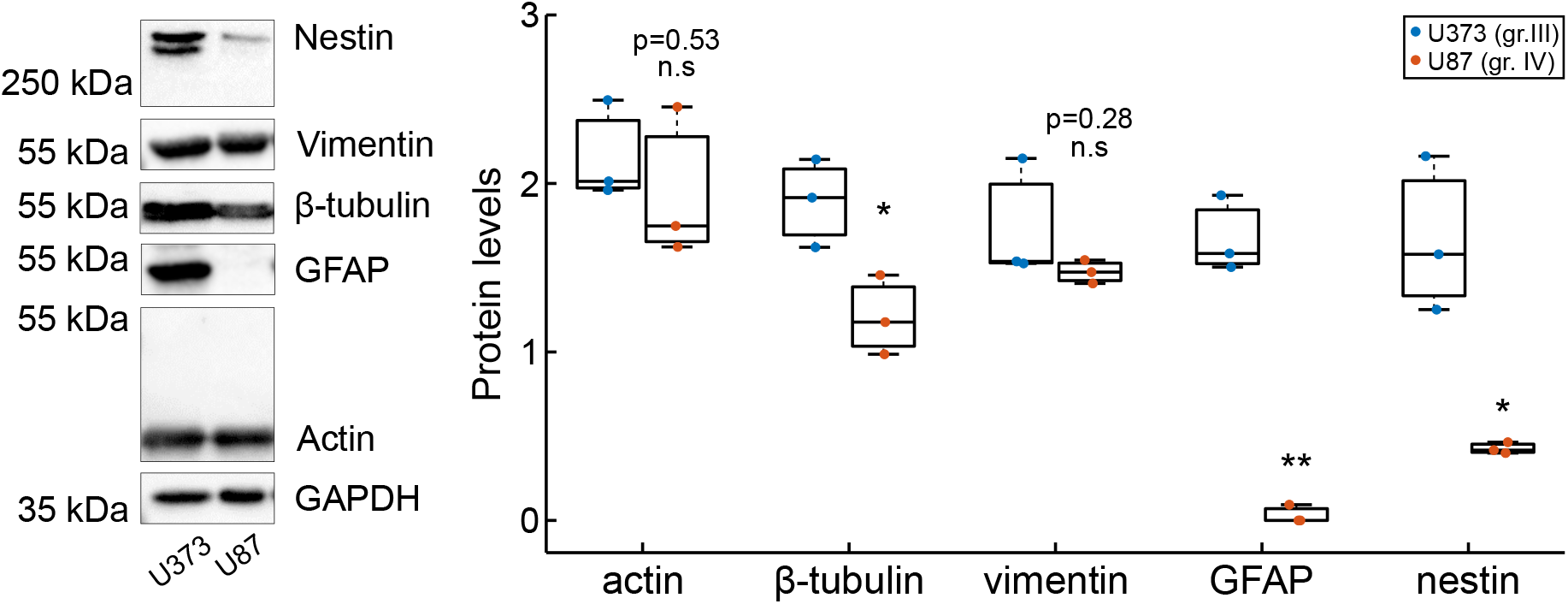
The expression of cytoskeletal proteins is lower in grade IV glioma. Expression levels of cytoskeletal proteins quantified by Western Blot in grade III (blue dots) and grade IV cells (red dots). GAPDH was used as a control for normalization. Results were obtained from at least three independent experiments.

### The spatial distribution of cytoskeletal fibers is consistent with an intracellular stiffening in grade IV gliomas

To get further insight into the role of cytoskeletal fibers in intracellular rheology, we imaged the organization of F-actin, microtubules and vimentin, GFAP and nestin intermediate filaments in cells plated on crossbow-shaped micropatterns (Fig. 4). Quantifying the levels of cytoskeletal fibers by immunofluorescence (Fig. 4A) confirmed the decrease in tubulin, GFAP and nestin measured by Western blots (Fig. 3) and also showed a statistically significant decrease in F-actin and vimentin staining in grade IV cells (Fig. 4A). However, in grade IV cells, we found that F-actin formed very short filaments dispersed in the perinuclear region in contrast with longer cortical stress fibers in grade III cells (Fig. 4A,B). 65.6% of grade IV cells (21 out of 32 cells) compared to 25.5% of grade III cells (14 out of 55 cells) displayed short perinuclear actin filaments. Intermediate filaments were more densely packed in the perinuclear region in grade IV cells and the cage formed by vimentin intermediate filaments around the nucleus, which was previously described in other cell types (Patteson et al., 2019), was more visible in grade IV cells (Fig. 4A,C). 75.4% of grade IV cells (43 out of 57 cells) compared to 26.2% of grade III cells (17 out of 65 cells) displayed a clear vimentin cage. Since intracellular microrheology essentially probes the perinuclear region (Fig. S4F), dispersed cytoplasmic short actin filaments and denser perinuclear intermediate filament networks may account for the increased intracellular rigidity in grade IV cells.

**Figure 4.**
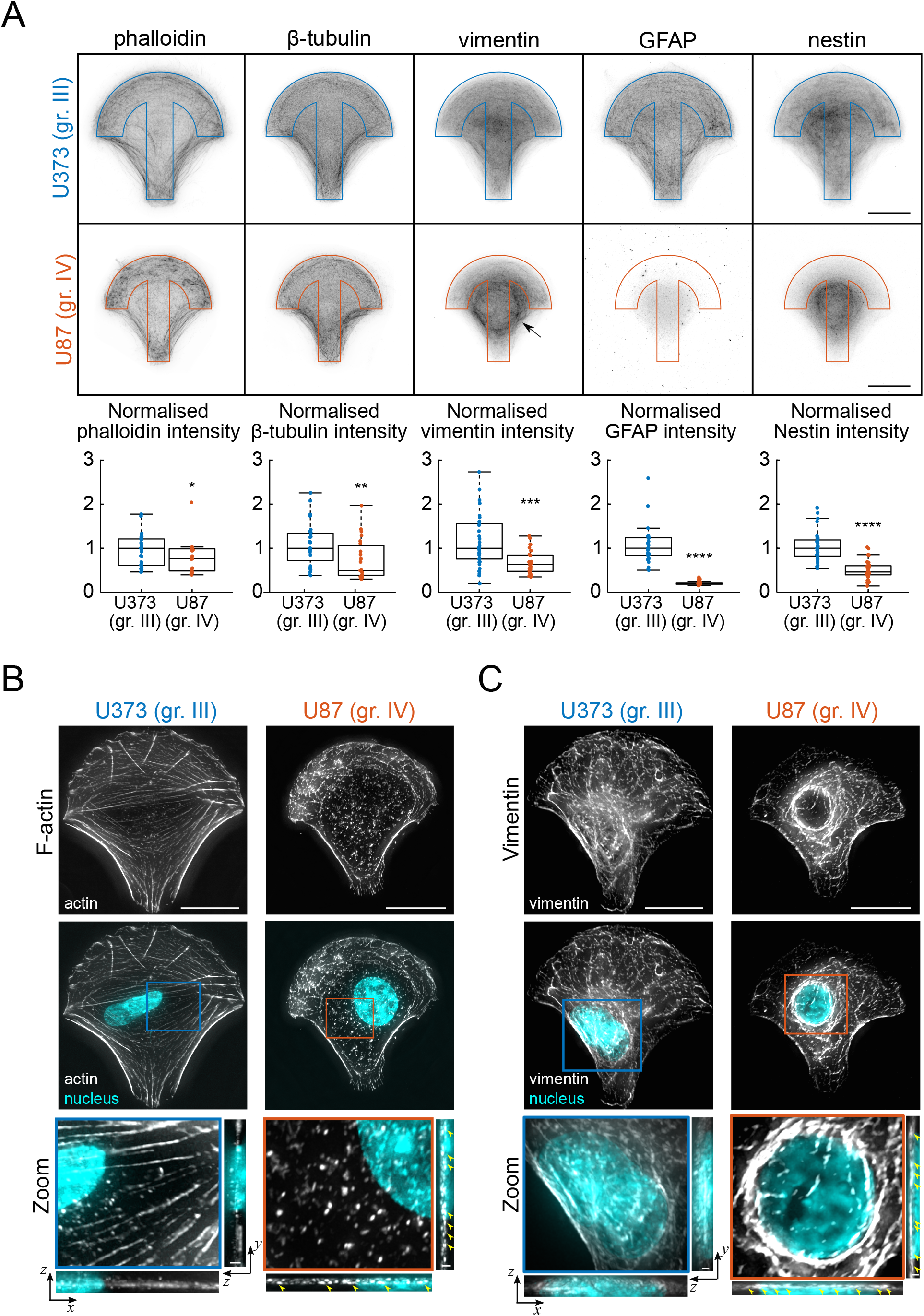
Spatial distribution of cytoskeletal networks in grade III and grade IV glioma cells plated on adhesive micropatterns. A. Averaged maps of the fluorescence intensity of F-actin (phalloidin), microtubules (β-tubulin) and intermediate filaments (vimentin, GFAP, nestin). Images are averages of n=55/32, 62/39, 65/57, 25/33, 30/33 cells grade III/IV cells for actin, microtubule, vimentin, GFAP and nestin intermediate filaments, respectively. The arrow on the vimentin map points to the increased density in perinuclear intermediate filaments in grade IV cells. Scale bar: 20 μm. The lower graphs show the fluorescence intensity levels of F-actin stained with phalloidin and β-tubulin, vimentin, GFAP and nestin stained with their corresponding antibodies. Data were obtained from n=30/17, 32/23, 38/29, 34/35, 35/38 grade III/IV cells for actin, tubulin, vimentin, GFAP and nestin, respectively. B-C. Typical examples of the organization of F-actin (B) and vimentin (C) filaments in U373 (grade III) and U87 (grade IV) glioma cells. F-actin and vimentin filaments are shown in grey scale and the nucleus in cyan. Zooms in the perinuclear region (boxed regions) are shown under the images. The *x-z* and *y-z* maximum intensity projections are shown below and to the right of the zoom images respectively. Arrowheads point to short actin filaments and perinuclear vimentin intermediate filaments surrounding the nucleus in U87 grade IV cells. Scale bars, 20 μm for the upper images and 2 μm for *x-z* and *y-z* projections in the zoom images.

### The nucleus is stiffer in grade IV than in grade III glioma cells

Since internal membranes were shown to contribute to intracellular stiffness (Mandal et al., 2016) and the nucleus is a major player in mechanotransduction (Thorpe and Lee, 2017), intracellular organelles could contribute to the mechanical differences we observed between grade III and grade IV glioma cells. We first asked whether the organization of intracellular organelles could be different in grade III and grade IV cells plated on micropatterns. Averaged immunofluorescence maps of the Golgi apparatus and of lysosomes indicate that these organelles have a wider distribution around the perinuclear region, the region probed by the intracellular method, in grade IV cells (Fig. S6A,B and S4F). The full width at half maximum of the normalized averaged fluorescent signal was 1.42 and 1.29 times larger in grade IV cells for Golgi and lysosomal membranes respectively (0.368 and 0.524 for the intensity profile of Golgi membranes in grade III and grade IV cells respectively, and 0.543 and 0.701 for the intensity profile of lysosomal membranes in grade III and grade IV cells respectively) suggesting that the density of Golgi and lysosomal membranes is larger in grade IV cells. The nucleus was shifted backwards in grade III cells, while it remained at the center of the micropattern in grade IV cells, suggesting that cell polarity is lost in grade IV glioblastoma cells (Fig. S6A,B) as reported previously (Camand et al., 2012). Quantification of the polarity of the nucleus-centrosome axis showed that it was indeed lower in grade IV cells (Fig. S6C), suggesting that a lower level of intracellular organization could be responsible for the wider average distribution of intracellular membranes. A higher probability of presence of internal membranes, such as Golgi or lysosomal membranes, in the perinuclear region could contribute to the increase in intracellular rigidity measured in grade IV cells.

Finally we asked whether the mechanical properties of the nucleus, which is a main contributor to cell stiffness (Caille et al., 2002; Mathieu and Manneville, 2019), could participate in glioma cells rigidity. Comparing the morphology of the nucleus in both glioma cell types, we found that the nucleus of grade IV cells is smaller, with a 2.2-fold smaller projected area (Fig. S7A,B), and more spherical (Fig. S7C) than that of grade III cells, consistent with the averaged maps and intensity profiles of the nucleus shown in Figure S6A,B. Taken together, these morphological data suggest that nuclear mechanics may differ between the two cell lines.

To quantify the mechanical properties of the nucleus, we performed indentation experiments using a bead held by the optical tweezers. An internalized bead initially in contact with the nucleus (visualized with Hoechst) was selected and trapped in the center of the optical tweezers. The nucleus was then pushed against the bead by moving the stage at a constant velocity of 42 nm/s (corresponding to a 2.5 μm displacement in 1 min) (Fig. 5A). The bead induced a nuclear deformation indicative of the rigidity of the nucleus. When the resistance of the nucleus became too strong, the bead was ejected from the trap. The shape of the nucleus then relaxed, at least partially, towards its initial shape (Fig. 5B). The time at which ejection from the trap occurred was significantly shorter (Fig. 5B, right) and the indentation of the nucleus was less pronounced in grade IV cells, indicating that the nucleus is stiffer in this cell type (Fig. 5C). To get more quantitative information regarding the mechanics of the nucleus, we analyzed the indentation phase using a simple Kelvin-Voigt viscoelastic model (Materials and Methods) that yields the nucleus spring constant *K*_*n*_ and the nucleus viscosity *η*_*n*_. The results show a statistically significant increase in elasticity *K*_*n*_ in grade IV cells (Fig. 5D). An increase in viscosity *η*_*n*_ was also observed but was not statistically significant (p=0.16).

**Figure 5.**
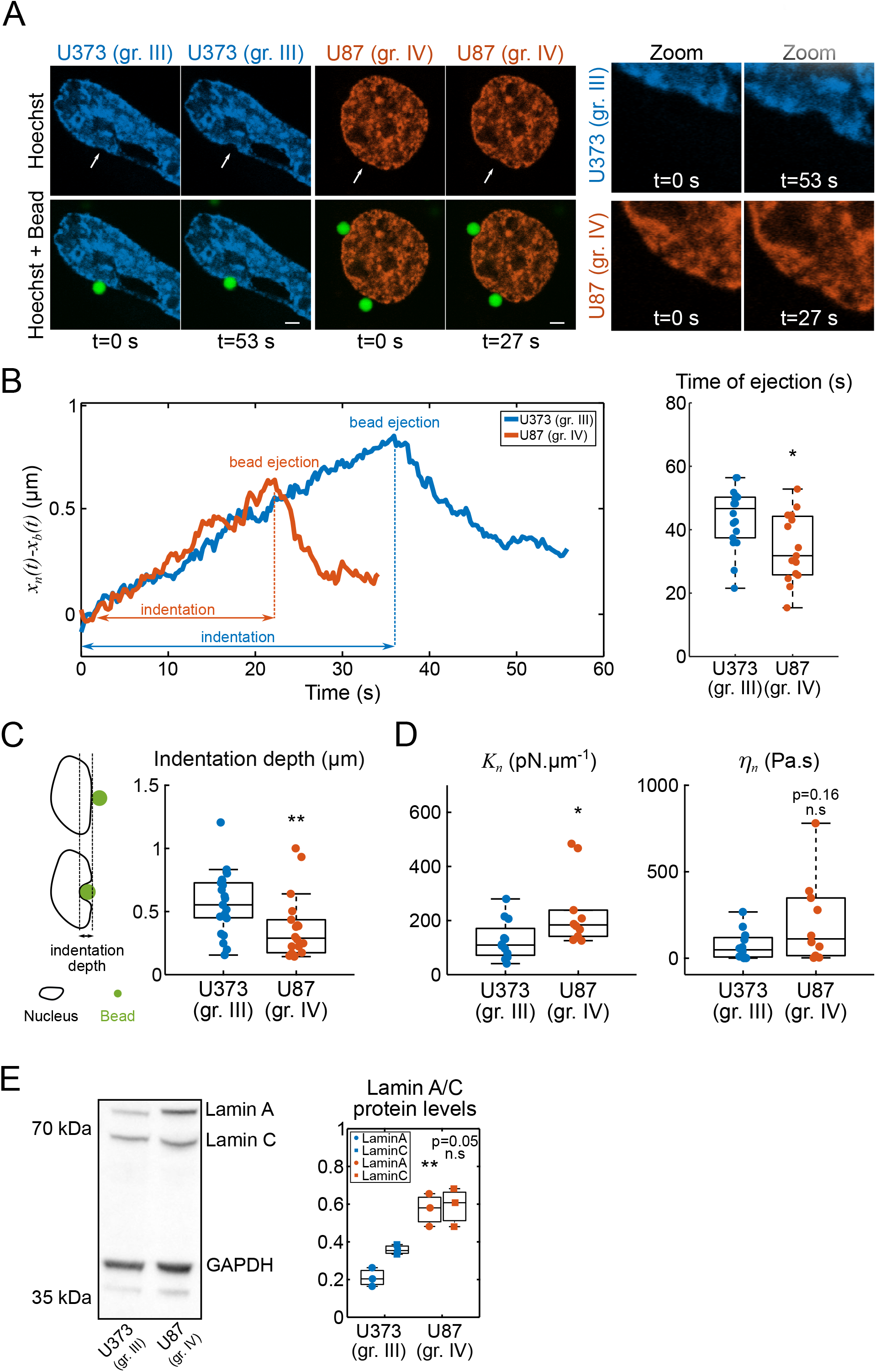
The nucleus is softer in grade III than in grade IV glioma cells. A. Typical nucleus indentation experiments in U373 grade III (blue) and U87 grade IV (orange) cells. Confocal images corresponding to Supplementary Movie S1. The nucleus was stained with Hoechst and the beads are shown in green. Scale bars, 2 μm. B. Typical plots of the nucleus and bead displacements as a function of time. Note that the plots were obtained from a different experiment than shown in Supplementary Movie S1 and A. The plot on the right shows the quantification of the time at which the bead is ejected out of the optical trap. Data were obtained from n=18 and 15 grade III and grade IV cells, respectively. C. Quantification of the maximal indentation depth. Data were obtained from n=23 and 18 grade III and grade IV cells, respectively. D. Rheological measurements of the nucleus from a viscoelastic analysis of the indentation phase (see Materials and Methods).The spring constant *K*_*n*_ and viscosity *η*_*n*_ of the nucleus were obtained by fitting the relaxation curves (grey lines) with a viscoelastic (Kelvin-Voigt) model. Data were obtained from n=12 and 10 grade III and grade IV cells, respectively. E. Lamins A/C expression levels are higher in U87 grade IV glioma cells than in U373 grade III cells. Typical Western blot (left) and corresponding quantification (right) form N=3 independent experiments.

Since lamins have been shown to contribute to the rigidity of the nucleus (Davidson et al., 2014; Harada et al., 2014; Lammerding et al., 2004; Liu et al., 2014), we have compared the levels of lamins A/C in grade III and grade IV glioma cells. Quantification by Western blots showed an increase in the expression of lamins A/C in grade IV cells (Fig. 5E), which is likely responsible for the increased rigidity of their nucleus.

## Discussion

We show here that a combination of whole cell and intracellular rheology techniques can efficiently discriminate the mechanical properties of glioma cells of different grades. Rat glioblastoma F98 cells are softer than normal primary rat astrocytes using both techniques. In contrast, human grade IV U87 cells are softer than grade III U373 cells when probed with the uniaxial single cell parallel plates rheometer, but stiffer when probed with intracellular optical-tweezers rheology. The spatial distribution of cytoskeletal fibers and internal compartments as well as nuclear mechanics may contribute to the observed mechanical differences.

Whole cell microrheology and intracellular microrheology do not probe the same cell compartments (Wu et al., 2018) and allow us to more precisely characterize cell mechanics. Whole cell rheology techniques, such as the uniaxial single cell parallel plate rheometer we used in this study (Bufi et al., 2015a), microfluidic-based devices (Darling and Di Carlo, 2015; Otto et al., 2015), the optical stretcher (Guck et al., 2005) or AFM using large probe tips (Grady et al., 2016), integrate the mechanical response of several cellular components and do not distinguish the contribution of individual cellular elements unless the cells are modified pharmacologically or genetically. It is generally assumed that both the nucleus and the actin cortex contribute to whole cell mechanical measurements (Caille et al., 2002). In our single cell parallel plate rheometer set-up, given the amplitude of the microplate displacement (typically 1μm, Fig. S2B), the measurements should be dominated by the actin cortex and not the nucleus, as is likely the case for real-time deformability cytometry (Urbanska et al., 2018). In contrast, intracellular rheology probes mechanics locally and is sensitive to the internal spatial organization of the cell. We observed that standardizing the intracellular organization is critical to detect subtle mechanical differences between grade III and grade IV glioma cells at the intracellular scale (Fig. 2 and S3). Fibronectin-coated adhesive micropatterns adapted to the size of the cells (i.e. their volume and spreading area) impose the same aspect ratio for both cell lines and avoid geometrical artifacts (Fig. S4). In our experimental conditions, the beads used for intracellular rheological measurements are embedded at least 1 μm deep within the cytoplasm (Guet et al., 2014), ruling out potential frictional effects from the actin cortex. Changes in signaling affecting intracellular mechanics may have local effects that are not measurable at a global scale. Because probing cell mechanics at different scales give complementary information, a multi-scale approach would be required for the development of a diagnostic tool.

Whole cell and intracellular rheology allowed us to compare the values of several mechanical parameters in grade III and grade IV glioma cells: the complex modulus *G*, phenomenological parameters (bead step displacement *X*_*b*_and rigidity index *RI*) and viscoelastic values (spring constant *K* and viscosity *η*). The absolute values of these parameters depend on a calibration step (see Materials and Methods) and on a characteristic length scale for stress propagation (Mandal et al., 2019, 2016). Furthermore, it was shown recently using breast cancer cells that mechanical parameters measured by different techniques and models can vary over several orders of magnitude (Wu et al., 2018). We thus focused this study on the mechanical differences between the two grades of glioma cells more than on the absolute values of the measured parameters. Our main finding is that the U87 grade IV glioblastoma cell line has a softer cortex but a stiffer intracellular compartment than U373 grade III astrocytoma cells (Fig. 2). We rationalize these opposite trends between cortical softening and intracellular stiffening by showing differences in cytoskeletal organization (Fig. 3 and 4), positioning of intracellular organelles (Fig. S6) and nuclear deformability (Fig. 5). Despite reduced levels in grade IV cells of tubulin and of the two intermediate filament proteins GFAP and nestin, two classical markers of glioblastoma (Quick et al., 2015) (Fig. 3), the perinuclear cytoplasm is stiffer in grade IV cells plated on micropatterns. This is most probably due to the spatial organization of the cytoskeleton. In cells plated on micropatterns, actin forms disorganized short perinuclear filaments in grade IV cells that may contribute to stiffen the cytoplasm (Fig. 4 and 6). More importantly vimentin intermediate filaments appear to form a dense cage around the nucleus of grade IV cells, as reported in other cell types (Mandal et al., 2016; Patteson et al., 2019), which is not as apparent in grade III cells (Fig. 4 and 6). Internal membrane compartments also appear to contribute to the intracellular stiffening in grade IV glioblastoma cells. We found that U87 grade IV cells plated on crossbow shaped micropatterns do not exhibit front-rear polarity, as opposed to U373 grade III cells (Fig. S6), in agreement with results obtained in migrating cells (Camand et al., 2012). Loss of polarity in grade IV cells may induce a higher average density of intracellular compartments around the nucleus, which could locally increase cytoplasmic stiffness. Finally, our indentation experiments show that the nucleus of grade IV cells is stiffer than that of grade III cells, consistent with increased lamin A/C levels in grade IV cells (Fig. 5), suggesting that nuclear stiffness contributes to intracellular mechanics. The dense vimentin filament network surrounding the nucleus may also contribute to our measurement of nuclear stiffness. To minimize the contribution of vimentin intermediate filaments in indentation experiments, we selected beads that were in close apposition to the nucleus when visualized by Hoechst staining (Fig. 5A, Movie S1). Furthermore, U87 grade IV cells are hypodiploid cells while U373 cells are near triploid cells (Conde et al., 2017), suggesting that our optical-tweezers nuclear indentation method is sensitive mostly to the mechanics of the nuclear envelop and less to the mechanics of chromatin.

**Figure 6.**
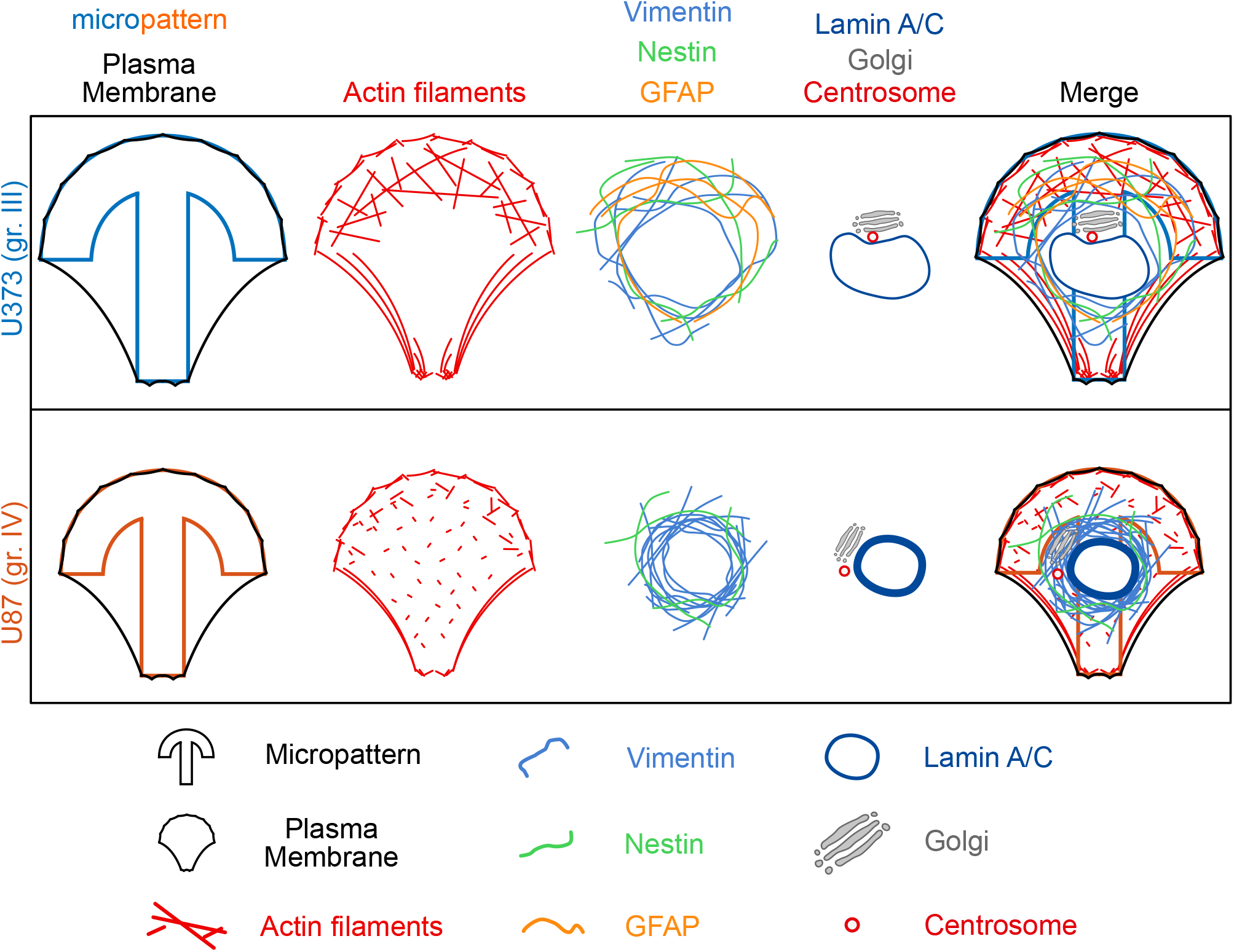
Scheme recapitulating the putative cellular elements contributing to the rheological differences between U373 grade III (top) and U87 grade IV (bottom) glioma cells.

It was shown that forces can propagate through the cytoplasm on distances of several micrometers from the cell surface to the nucleus (Poh et al., 2012; Tajik et al., 2016). Our results suggest that, conversely, the stiffness of the nucleus can impact the stiffness of the cytoplasm when measured by the viscoelastic relaxation of a bead located several micrometers away from the nucleus. In contrast, measurements with the single cell microplate rheometer, which report a softening of grade IV cells, do not seem to be influenced by nuclear stiffness.

Most studies so far, including ours in breast cancer cells (Mandal et al., 2016), show that individual cancer cells soften as their metastatic potential increases (Alibert et al., 2017). Such softening could increase cell deformability and facilitate the migration of cancer cells from the primary tumor and their invasion into the surrounding tissues during the metastatic process. Since the nucleus is thought to be the limiting factor when a cell migrates through confined spaces, for instance through narrow pores or constrictions *in vitro* (Denais et al., 2016; Lautscham et al., 2015; Raab et al., 2016), the nucleus may also be softer in cancer cells, in correlation with decreased levels of lamins A/C (Pajerowski et al., 2007). In agreement with this hypothesis, micropipette aspiration experiments show an increase in nuclear deformability in epithelial-to-mesenchymal transition (EMT)-induced mammary epithelial tumor cells (Mekhdjian et al., 2017). In this study however, as in studies based on cells migrating in microfluidic devices (Denais et al., 2016; Lautscham et al., 2015; Raab et al., 2016) or based on real-time deformability cytometry (Holenstein et al., 2019), the mechanics of the nucleus is not probed directly and the actin cortex may also participate in the observed softening. By measuring directly nuclear stiffness in living cells (Fig. 5), we show here that in the case of grade IV U87 glioblastoma cells, the nucleus is stiffer than in grade III U373 cells despite cortical softening. Consistently, grade IV cells exhibit elevated levels of lamins A/C compared to grade III cells. Glioblastomas are generally softer than the surrounding brain parenchyma (Streitberger et al., 2014). The extremely soft microenvironment of glioblastomas and the complex composition of the extracellular matrix in vivo (Pogoda et al., 2017) may be related to the atypical mechanics of glioblastoma cells. The rigidity of the nucleus may not be a hindrance to cell invasion in the soft brain tissue and could explain why glioblastomas do not extravasate and metastasize. The behavior we described here may not be typical of all glioblastoma cells but shows that assessing the mechanics of primary glioblastoma cells derived from patients at different scales will be critical to develop reliable diagnostic and/or prognostic methods based on mechanophenotyping.

## Materials and Methods

### Cell Culture

Human glioma cell lines U373 (grade III astrocytoma, also distributed as U251 cells) and U87 (grade IV glioblastoma) were grown in MEM-GlutaMAX medium (Gibco) supplemented with 10% FCS (Eurobio) and 1% Non-Essential Amino Acids (Gibco). Primary astrocytes were obtained from E17 rat embryos following (Etienne-Manneville, 2006). They were grown in DMEM supplemented with 10% FCS (Eurobio). Rat glioblastoma cell lines F98 were cultured in DMEM-F12 (Gibco) supplemented with 10% FCS (Eurobio). All cells were cultured at 37°C with 5% CO_2_.

### Single Cell Microplate Whole Cell Microrheology

Whole cell microrheology experiments were performed using a set-up described previously (Bufi et al., 2015a; Desprat et al., 2006) (Fig. S1A). Briefly, primary astrocytes and glioma cells were caught between one rigid and one flexible glass microplates, previously coated with a 2% pluronic F127 solution (Sigma-Aldrich) for 45 min. Oscillations in the frequency range of 0.1–6.4 Hz were applied to the flexible plate by a computer-controlled piezoelectric stage. The deflection of the flexible plate (of calibrated stiffness) was monitored by an optical sensor, allowing us to measure the stress *σ*(*t*) applied to the cell and the resulting strain *ɛ*(*t*). The compression modulus *E*(*ω*) is defined from the ratio *σ*(*t*)/*ɛ*(*t*) and can be expressed as *E*(*ω*) = *E*′(*ω*) + *iE*″(*ω*), where *E*′(*ω*) and *E*″(*ω*) are the storage and the loss moduli. Knowing the amplitude and the phase of the two signals **σ**(*t*) and *ɛ*(*t*), we can deduce the values of *E*′ and *E*″ for each frequency. In this frequency range, cells were shown to exhibit a power-law behavior *E*(*ω*) = *E*_0_*ω*^α^. The shear modulus *G* can be obtained from the elongation modulus *E* using 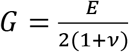 where *v* is the Poisson coefficient. Since our aim was to compare the characteristic moduli of cells at different stages of cancer progression, we used for simplicity simple power laws to fit the *G*′ and *G*″ data over the whole frequency range 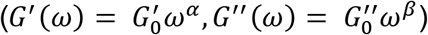, even if the *G*″ data suggested a more complex behaviour with an additional viscous-like component, the effect of which is visible at high frequencies (Fig. 1A), in agreement with the damping model introduced previously by Fabry, Fredberg and colleagues (Fabry et al., 2001). The amplitude of the deformation was kept in the limit of linear elasticity solicitations (10–15% of the cell size).

### Optical Tweezers-Based Intracellular Microrheology

The setup combining optical tweezers and fast confocal microscopy was described in detail previously (Guet et al., 2014). Briefly, a single fixed optical trap was built by coupling a 1060-1100 nm infrared laser beam (2W maximal output power; IPG Photonics) to the back port of an inverted Eclipse microscope (Nikon) equipped with a resonant laser confocal A1R scanner (Nikon), a 37°C incubator, and a nanometric piezostage (Mad City Labs). Coverslips with micropatterned cells containing typically one to three internalized beads were mounted in a Ludin chamber. The piezostage was displaced either in a stepwise fashion for step relaxation experiments or in a periodic fashion for oscillatory microrheology experiments.

Step relaxation experiments (Fig. S1B) were described previously (Guet et al., 2014). Briefly, a 2 μm-diameter bead was first trapped by the fixed optical tweezers (1W laser output power, corresponding to 150 mW on the sample, trap stiffness *k*_*trap*_ between 200 and 280 pN.μm^−1^) calibrated by the Stokes drag force method). Following a *X*_*s*_ = 0.5 μm step (linear increase in 40 ms) displacement of the stage, the bead was moved out of the trap center due to the viscoelasticity of its microenvironment. As the optical tweezers acts as a spring on the bead, the bead position *x*_*b*_(*t*) relaxes from its maximal position *X*_*b*_, termed bead step amplitude, toward the center of the optical trap. Relaxation curves *x*_*b*_(*t*) during a duration *T* = 10 s are obtained with a home-made Matlab tracking code and analyzed with three methods : a phenomenological approach, a power-law based analysis and a viscoelastic standard linear liquid (SLL) model analysis as described previously (Mandal et al., 2019, 2016). The phenomenological approach gives the bead step amplitude *X*_*b*_ and the rigidity index *RI* defined as 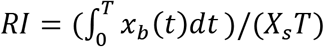 so that *RI* → 0 corresponds to a very soft and deformable cytoplasm while *RI* → 1 corresponds to a very stiff and rigid cytoplasm. The power law analysis yields the complex shear modulus *G* = *G*′ + *iG*″ with *G*′ the storage modulus (elastic-like behaviour) and *G*″ the loss modulus (viscous-like behaviour). The amplitude of the complex shear modulus is given by 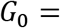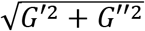 The SLL model yields the spring constant *K* and the viscosity *η*.

Oscillatory intracellular rheology (Fig. S1C) was performed by applying a *X*_*s*_ = 0.5 μm-amplitude sinusoidal displacement at 0.8 Hz for 40 sec of the stage. The displacements of a bead trapped in the optical tweezers (1 W laser output power, corresponding to 150 mW on the sample) and of a reference bead located in the cytoplasm away from the optical trap were recorded at 120 frames/s in the resonant mode of the confocal scanner for 40 s using the NIS Nikon software. The displacement of the trapped bead relative to the reference bead was fitted as in (Mandal et al., 2019) to obtain the elastic storage modulus *G*′ and the viscous storage modulus *G*″.

### Morphological characterization of glioma grade III and IV cells

The diameter of glioma cells was determined from cells ins suspension using Differential Interference Contrast (DIC) microscopy and image analysis with ImageJ. Cell volume was deduced from the measurements of the diameter of non-adhering cells assuming a spherical geometry (*V* = 4/3*πR*^3^). We found *V* = 4198 ± 1555 μm^3^ and *V* = 2909 ± 191 μm^3^ (median values ± standard error mean) for U373 (grade III) and U87 (grade IV) cells respectively. To determine the surface area of glioma cells, cells were plated either on glass or on fibronectin, incubated with the cell marker Cell Mask (Invitrogen) for 20 minutes, washed with PBS, and imaged right away. Fluorescence images of the bottom plane of the cells were taken and thresholded with ImageJ to determine the surface area. We found *S*_*glass*_ = 1988 ± 131 μm^2^, *S*_*glass+FN*_ = 2095 ± 121 μm^2^ and *S*_*glass*_ = 1427 ± 117 μm^2^, *S*_*glass+FN*_ = 1390 ± 95 μm^2^(median values ± standard error mean) for U373 (grade III) and U87 (grade IV) cells respectively.

### Micropatterning

Micropatterns were printed by deep UV photolithography technique. 18-μm radius glass coverslips were plasma-cleaned for 5 min and incubated immediately with 0.1 mg/ml PLL-PEG diluted in HEPES 10 mM for 1h at room temperature. Coverslips were washed twice in PBS and dried before printing. Micropatterns were printed for 5 min with specifically designed chrome masks and coated for 1h at 5% CO_2_ and 37°C with 50μg/mL fibronectin and 20μg/mL Alexa 546–fibrinogen (Sigma) diluted in distilled water. Micropatterned coverslips were washed twice in PBS and fresh medium. Cells were seeded on freshly prepared protein-coated micropatterns and allowed to spread for at least 2–3 h before experiment or fixation. Non-adherent cells were washed off by rinsing with culture medium. Crossbow patterns have a diameter of 50 μm and 61 μm. For rheological measurements, cells were incubated overnight with 2-μm-diameter fluorescently labeled latex beads (580/605 or 365/415 fluorescence; Bangs Laboratories) before seeding them on the micropatterns. 20 mM Hepes was added to the medium before microrheology experiments.

### Immunofluorescence, Western Blot and reagents

For immunofluorescence experiments, cells were fixed with 4% paraformaldehyde for 15 minutes at room temperature or with cold methanol for 3 minutes at −20°C. Cells were then permeabilized and block with a mix of PBS+2% BSA+Saponin or PBS+2% BSA for 20 min at room temperature. Primary and secondary antibodies were diluted in PBS+ 2% BSA and incubated for 1h at room temperature, protected from light. Coverslips were finally mounted with Mowiol complemented with DAPI to visualize nuclei.

For Western blot experiments, cells lysates were obtained with Laemmli buffer composed of 120 mM Tris-HCl pH7.5, 4% glycerol, 4% SDS and 100 mM DTT. Samples were boiled 5 min at 95° before loading on 8% or 10% polyacrylamide gels. Migration and transfer occurred respectively at 150 V and 25 V. Membranes were blocked with PBS + 0.2% Tween and 5% milk during 1h and incubated 1h with the primary antibody and 1h with HRP-conjugated secondary antibody both diluted in PBS + 0.2% Tween. Bands were revealed with ECL chemoluminescent substrate (Thermo Scientific, SuperSignal West Femto).

Primary antibodies were anti-βtubulin (Sigma-Aldrich T4026), anti-vimentin (Santa Cruz sc-7557), anti-GFAP (Santa Cruz sc-6170), anti-nestin (Millipore ABD69 and MAB353). Anti-GM130 (BD Transduction Laboratories), anti-LAMP1 (Sigma-Aldrich L1418), anti-lamin (MANLAC1(4A7) from MDA Monoclonal Antibody Resource) (Manilal et al., 2004), and anti-GAPDH (Chemicon MAB374). The anti-γtubulin antibody was a gift from Michel Bornens. Actin was labeled using Alexa 546–phalloidin (Invitrogen) or anti-Actin (Sigma-Aldrich A4700). Secondary antibodies were from Jackson ImmunoResearch Laboratories. Nuclei or plasma membranes were stained in live cells after 20 min incubation with Hoechst (Sigma-Aldrich) or CellMask (Molecular Probes, Thermo Fisher) respectively.

### Quantifications of cytoskeleton proteins from immunofluorescence images

Images of fixed cells were acquired as confocal z-stacks separated by 0.2 μm in the galvanometric mode of the confocal scanner with a 4- or 8-frames averaging using the NIS-Elements Nikon software. Z-stacks were processed with a sum intensity projection using ImageJ. Cell contours were drawn manually and the mean fluorescence intensity inside the contour was measured.

### Intracellular spatial organization on micropatterns

Images of fixed cells were acquired as z-stacks separated by 0.2 μm with a Delta-Vision microscope using the Softworks software (objectives 60X or 100X). Images were deconvoluted using an enhanced-ratio 15. For each cell, the deconvoluted images were automatically aligned and processed with a maximum intensity projection using a custom-written ImageJ macro. The final image was obtained with an average z-stack projection of all the cells and displayed using an inverted look-up table (LUT). Averaged fluorescence profiles of intracellular organelles (Golgi apparatus, lysosomes, nucleus) were obtained by averaging the fluorescence intensity profiles from individual maximum intensity projections along a line aligned with the axis of the crossbow-shaped micropattern and of the same width as the micropattern (61 μm for U373 grade III cells and 50 μm for U87 grade IV cells). Images of nuclei were filtered with a Gaussian blur and thresholded with ImageJ to determine the nucleus area. An ellipse fit of the nucleus was used to determine the lengths of the major and minor axes and the aspect ratio of the nucleus.

### Nucleus indentation experiments using intracellular optical-tweezers based microrheology

The intracellular microrheology set-up was used to deform nuclei of different grades of glioma directly *in cellulo* and probe their mechanical properties. A bead, close to the nucleus, was first trapped with the optical tweezers. Then, the stage was displaced at a constant speed (2.5 μm in 1 min, *v*_0_ =41.7 nm/s) so that the nucleus was moved towards the bead and pushed against it. In a first phase, called indentation phase, the trap applied a spring force on the bead, keeping the bead inside the optical trap. As the nucleus moved against the bead, the bead indented the nucleus. The maximal indentation depth was measured by image analysis using ImageJ. When the force applied by the nucleus on the bead became larger than the spring force of the trap, the bead was ejected out of the trap and the nucleus relaxed towards its initial shape (relaxation phase) (see Supplementary Movie S1). The positions of the center of mass of the bead relative to its initial position *x*_*b*_(*t*) and of the center of mass of the nucleus relative to its initial position *x*_*n*_(*t*) (or of a small region representative of the global movement of the nucleus when the nucleus was not imaged entirely), were tracked using a home-made Matlab code. The spring constant *K*_*n*_ and viscosity *η*_*n*_ of the nucleus were obtained by fitting the relaxation of *x*_*n*_(*t*) − *x*_*b*_(*t*) with a viscoelastic (Kelvin-Voigt) model. Balancing the forces exerted on the bead of radius *R*, i.e. the force exerted by the trap and the viscoelastic forces exerted by the nucleus, gives 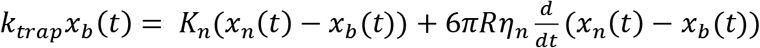 (Eq. 1). With *x*_*n*_(*t*) = *v*_0_*t* (linear displacement of the nucleus) and the initial condition *x*__*n*__(*t* = 0) − *x*_*b*_(*t* = 0) = 0, solving equation (Eq. 1) gives 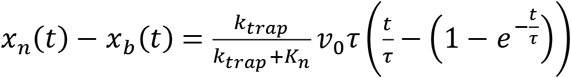 (Eq. 2) with 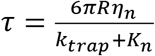 Fitting the curves (*x_n_* (*t*) − *x*_*b*_ (*t*)) with equation Eq. 2 yields the spring constant *K*_*n*_ and the viscosity *η*_*n*_ of the nucleus.

### Statistical analysis

All data are presented as the mean value +/- s.e.m. of at least three (N=3) independent experiments. Box plots show the median value, the first and third quartiles and whiskers correspond to ±2.7**σ** where **σ** is the standard deviation. Statistical relevance was evaluated using Student′s t-test or Wilcoxon rank test with Matlab, depending on the number of cells and the normality of the distribution. For experiments where n≥30 cells a Student′s t-test was used. When n<30 cells an Anderson-Darling test was used to evaluate normality. Data sets following a normal distribution were compared using a Student′s t-test, while non-parametric data sets were compared with a Wilcoxon rank test. p-values are reported as non-significant (n.s. p>0.05, p-values are indicated), or significant * p<0.05, ** p<0.01, *** p<0.001, **** p<0.0001.

## Acknowledgements

We thank Dalila Labiod for providing F98 cells. We thank Raffaële Attia for her help in photomask design. We thank Cécile Leduc, Kerren Murray and Chiara de Pascalis for help with primary rat astrocytes and transfection reagents. We thank Mathieu Coppey, Chloé Geller and Fiona Francis for stimulating discussions. We thank Sébastien Manneville, Sandra Lerouge and Sophie Asnacios for technical assistance. We thank Kalpana Mandal for technical assistance. C.A was funded by a 3 years grant from UPMC University Paris 6 (Programme Doctoral “Interfaces Pour le Vivant”) and a 6 months grant from La Ligue Contre le Cancer. Other fundings were from Institut Curie, CNRS, INSERM Plan Cancer 2009-2013 INSERM - CEA Tecsan (grant number PC201125), and the labex «Who AM I ?», labex ANR-11-LABX-0071 and the Université de Paris, Idex ANR-18-IDEX-0001 funded by the French Government through its «investments for the future» program.

## Supplementary figures legends

**Figure S1**

Microrheology techniques used to probe cortical and intracellular mechanics.

A. Uniaxial single cell microplate rheometer probing mechanics at the whole cell scale. Schematics depicting the applied displacement of the basis *D*(*t*) and the measured displacement of the tip *d*(*t*) of the flexible microplate.

B-C. Intracellular optical tweezers probing mechanics at the subcellular scale. Schemes depicting the relaxation (B) and oscillation (C) protocols.

**Figure S2**

Cortical rheology measured with the uniaxial single cell plate rheometer.

A. Schematics of the uniaxial single cell microplate rheometer experiments showing the applied displacement of the basis *D*(*t*) and the measured displacement of the tip *d*(*t*) of the flexible microplate in contact with a U373 grade III (blue) or a U87 grade IV (orange) glioma cell subjected to an oscillating stretch.

B. Typical examples of *D*(*t*) and *d*(*t*) for a U373 grade III (blue) or a U87 grade IV (orange) glioma cell subjected to an oscillating stretch at 0.4 Hz as a function of time. The sinusoidal fits of the experimental data are shown in straight lines.

**Figure S3**

Intracellular active microrheology cannot discriminate between freely adhering U373 grade III and U87 grade IV gliomas cells.

A. Averaged relaxation curves in grade III glioma (blue) and grade IV glioma (orange). Data were obtained from n=16 and 14 beads in grade III and grade IV cells, respectively.

B-C. Phenomenological parameters (bead step amplitude *X*_*b*_ and rigidity index *RI*) and parameters obtained by fitting the relaxation curves with the SLL model (spring constant K and viscosity *η*) (B) or with a power-law analysis (elastic storage modulus *G*′, viscous loss modulus *G*″) (C). Data were obtained from n=16 and 14 grade III and grade IV cells, respectively. p values were all above 0.2 indicating no statistical difference between grade III and grade IV cells.

D. Shear, storage and loss moduli measured by intracellular oscillatory microrheology in freely adhering grade III and grade IV glioma cells. Data were obtained from n=21 and 19 grade III and grade IV cells, respectively.

**Figure S4**

U373 grade III and U87 grade IV glioma cells have different sizes and different spread areas.

A-B. The spread area of U373 (grade III) cell is larger than that of U87 (grade IV) cells. A. Adhering grade III and grade IV glioma cells imaged by fluorescence microscopy (CellMask staining). Scale bar, 20 μm. B. Spread area of grade III and grade IV glioma cells on non-treated glass coverslips (Glass) or fibronectin (Glass+FN) coated glass coverslips. Data were obtained from n=38/58 and 41/50 (on glass/fibronectin) grade III and grade IV cells, respectively.

C-D. The volume of U373 (grade III) cell is larger than that of U87 (grade IV) cells. C. Non adherent grade III and grade IV glioma cells imaged with differential interference contrast (DIC) microscopy. Scale bar, 10 μm. D. Volume of grade III and grade IV glioma cells. Data were obtained from n=59 and 58 grade III and grade IV cells, respectively.

E. Schematics explaining the design of the adhesive micropatterns as a function of cell spread area and volume.

F. 2D maps of bead positions in grade III (left) and grade IV (right) glioma cells. Data were obtained from n=35 and 35 grade III and grade IV cells, respectively.

**Figure S5**

Typical examples of the bead displacement curves as a function of time in oscillatory microrheology experiments in U373 grade III and U87 grade IV glioma cells plated on adhesive micropatterns.

**Figure S6**

Spatial distribution of intracellular organelles in grade III and grade IV glioma cells plated on adhesive micropatterns.

A. Averaged maps of the fluorescence intensity of the Golgi apparatus (GM130), lysosomes (LAMP1) and the nucleus (DAPI) and superimposition of the positions of the centrosome (γ-tubulin). Images are averages of n=55/65, 25/33, 80/86, 30/32 grade III/IV cells for the Golgi apparatus, lysosomes, nucleus and centrosome stainings, respectively. Scale bar: 20 μm.

B. Averaged fluorescence profiles of the Golgi apparatus, lysosomes and nucleus staining along the axis of the crossbow-shaped micropattern in U373 grade III (blue lines) and U87 grade IV (orange lines) cells. The graphs show intensities normalized to the maximum value as a function of the distance *x* along the crossbow axis normalized by the size of the crossbow from the back (−1 < *x* < 0) to the front (1 > *x* > 0) of the pattern. The normalized distribution of the beads along the micropattern axis in both cell types (see also Fig. S4F) is superimposed as vertical bars on the normalized Golgi and lysosome intensity profiles to highlight that both organelles localize in the perinuclear region probed by the beads used in intracellular rheology experiments.

C. Polarization of the nucleus-centrosome axis. The position of the centrosome was scored as belonging to one of the four quadrants indicated in the schematics below the graph. Centrosomes located in the first quadrant indicate polarization towards the front edge of the crossbow.

**Figure S7**

Morphology and mechanics of the nucleus in grade III and grade IV glioma cells.

A. Typical images of the nucleus stained with DAPI in a grade III cell and in a grade IV cell plated on crossbow-shaped micropatterns (Scale bar, 20 μm).

B-C. Quantification of the nucleus projected area (B), of the lengths of the major and minor axis of an ellipse fit to the nucleus contour (C, left) and of the aspect ratio of the fitted ellipse (C, right). Data were obtained from n=80 and 86 grade III and grade IV cells, respectively.

## Supplementary movies

**Movie S1**(corresponding to Fig. 5A)

Nucleus indentation experiments in a U373 grade III astrocytoma cell (left) and in a U87 grade IV glioblastoma cell (right). Top movies show the 2μm-diameter bead in green and the nucleus stained by Hoechst in blue. Bottom movies only show the nucleus channel in grayscale. The cross indicates the position of the center of the optical trap. The arrow point to the nuclear deformation induced by the bead. Images from the movies are shown in Fig. 5A. The real-time duration of the movie is 2 min (115 frames at 1.04 sec/frames). The movie was accelerated 30 times for display. Scale bar, 5 μm.

