## Supplementary figures and images for "Intracellular stiffening competes with cortical softening in glioblastoma cells"

### Supplemental Figures

#### Figure S1

A

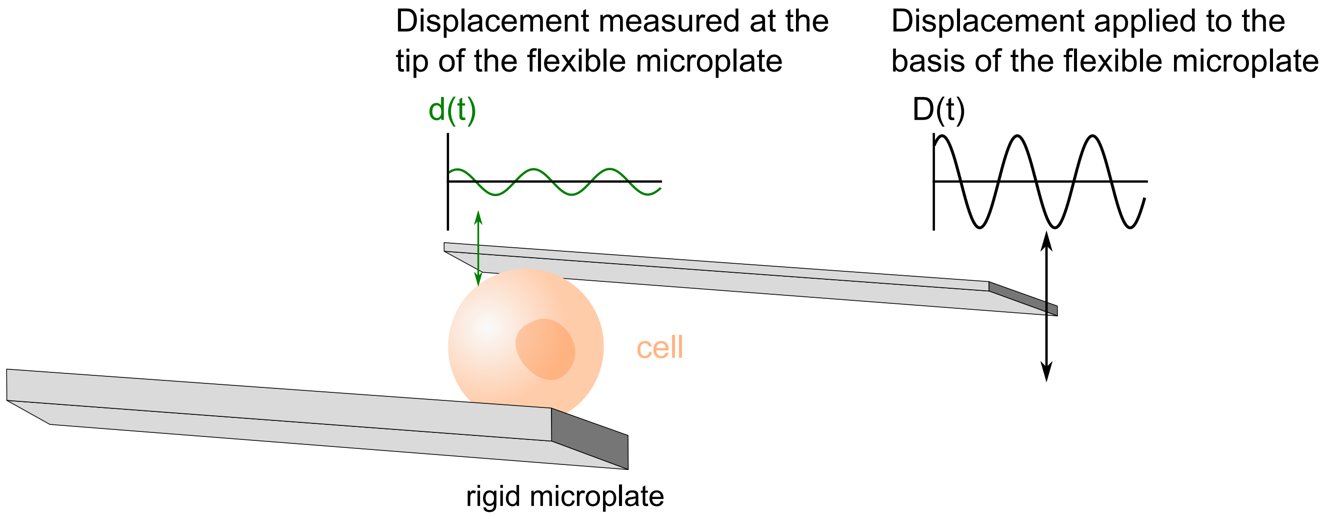


B

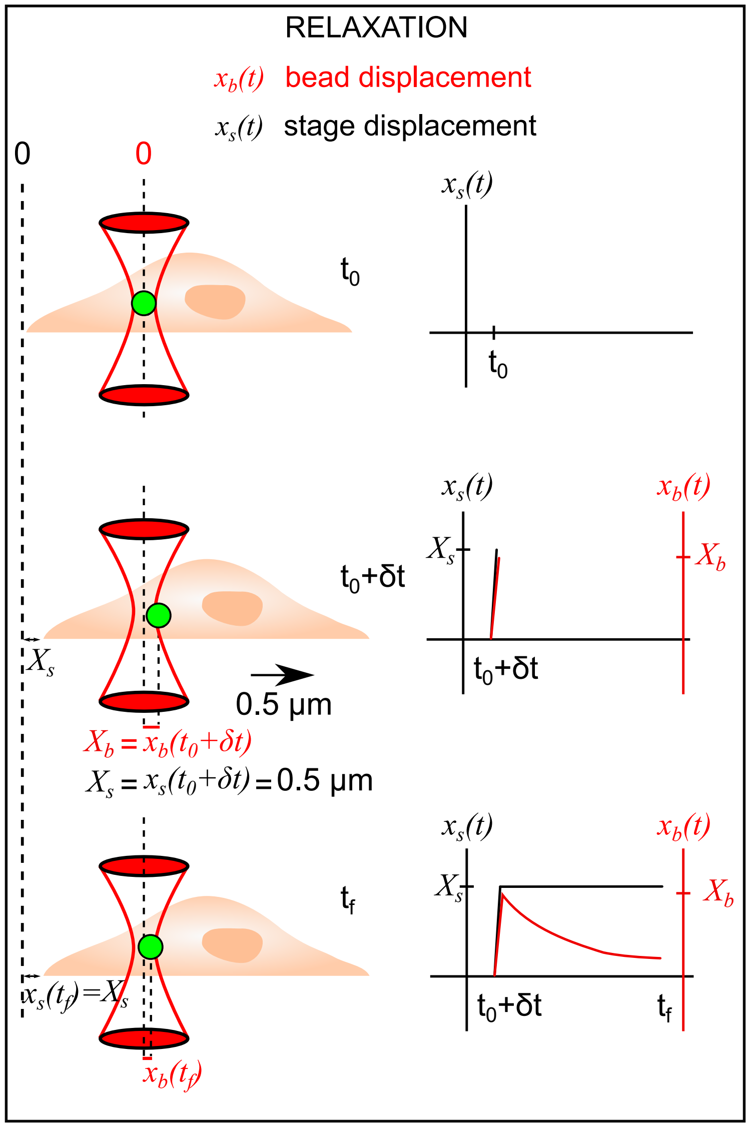


C

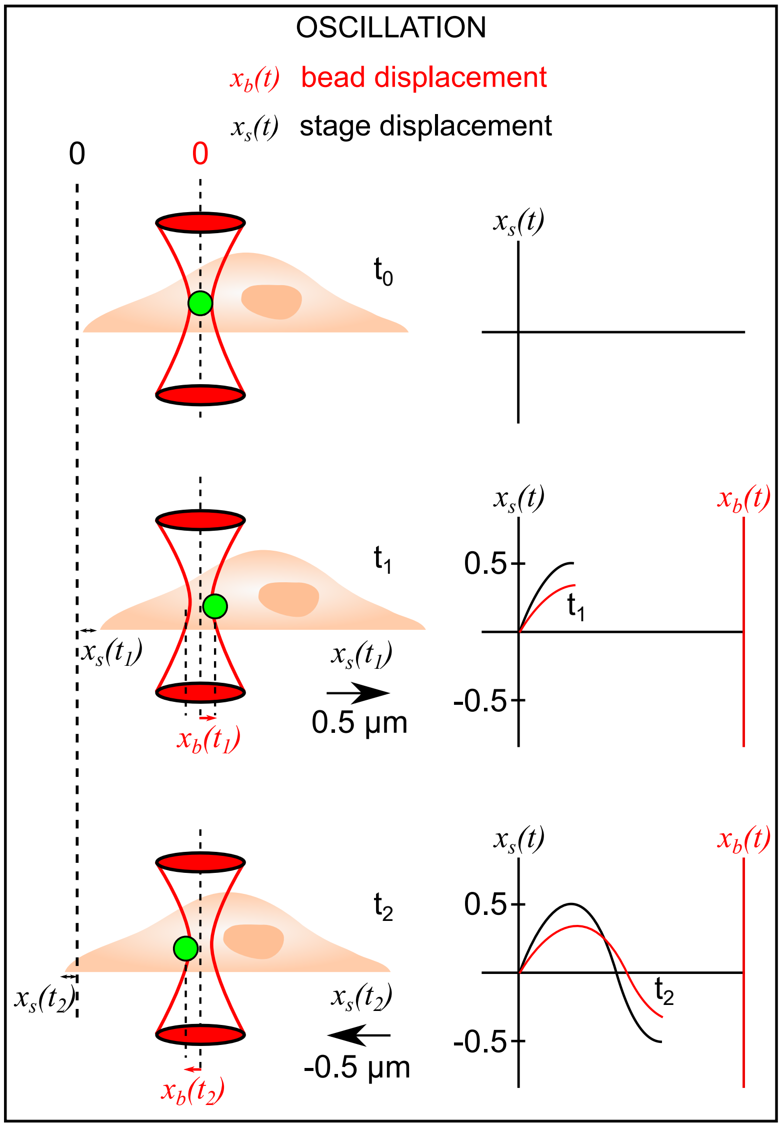

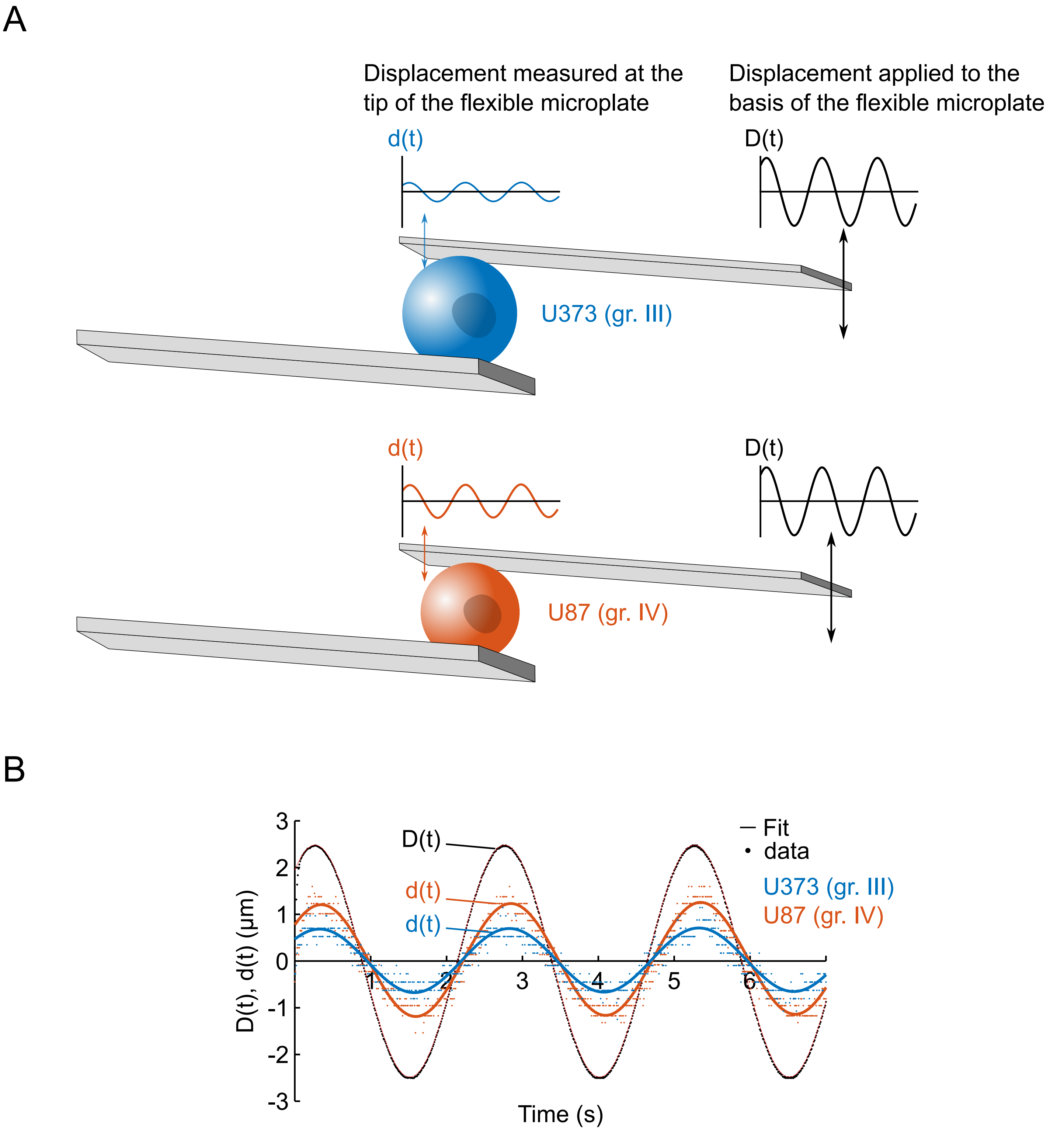

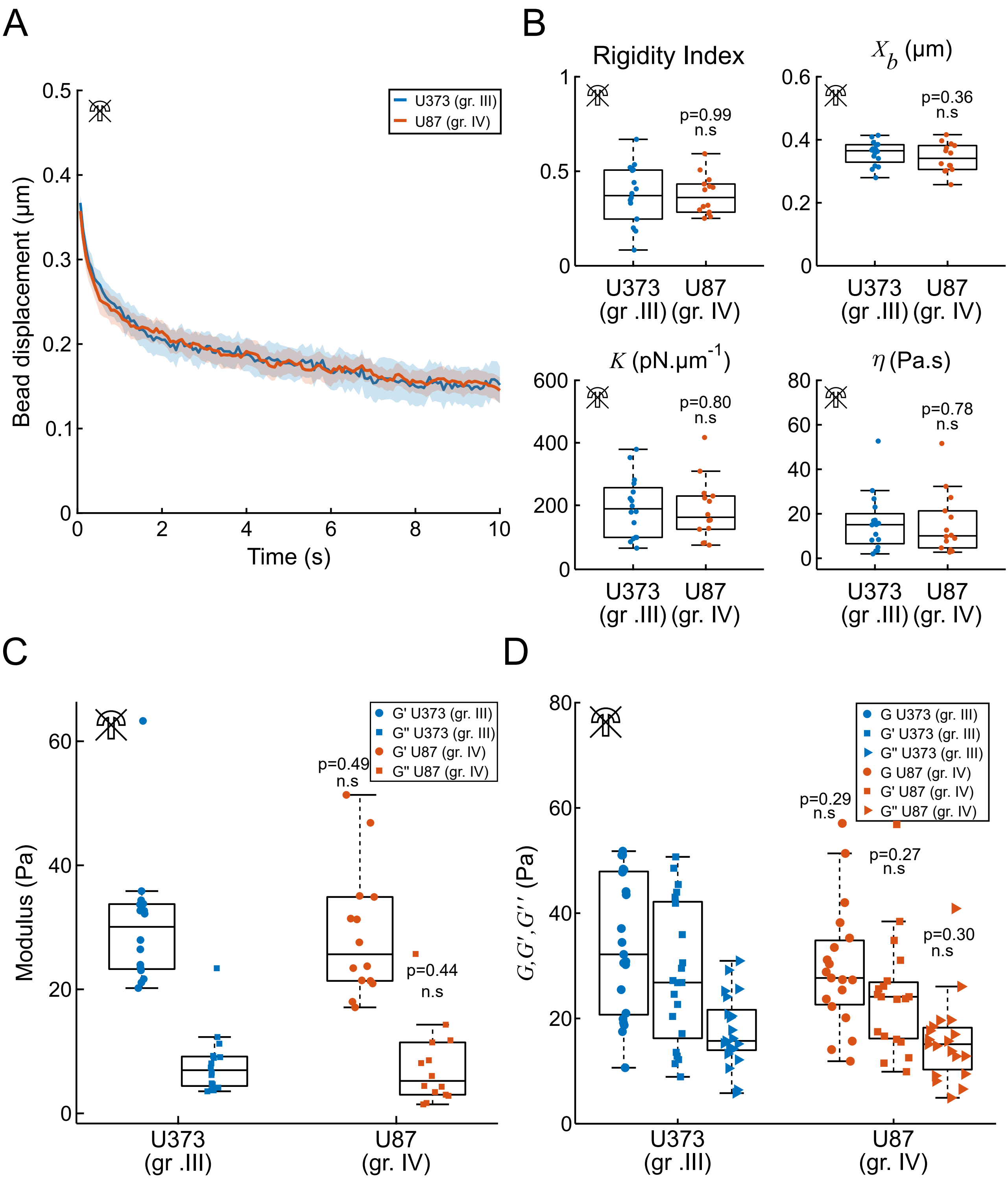

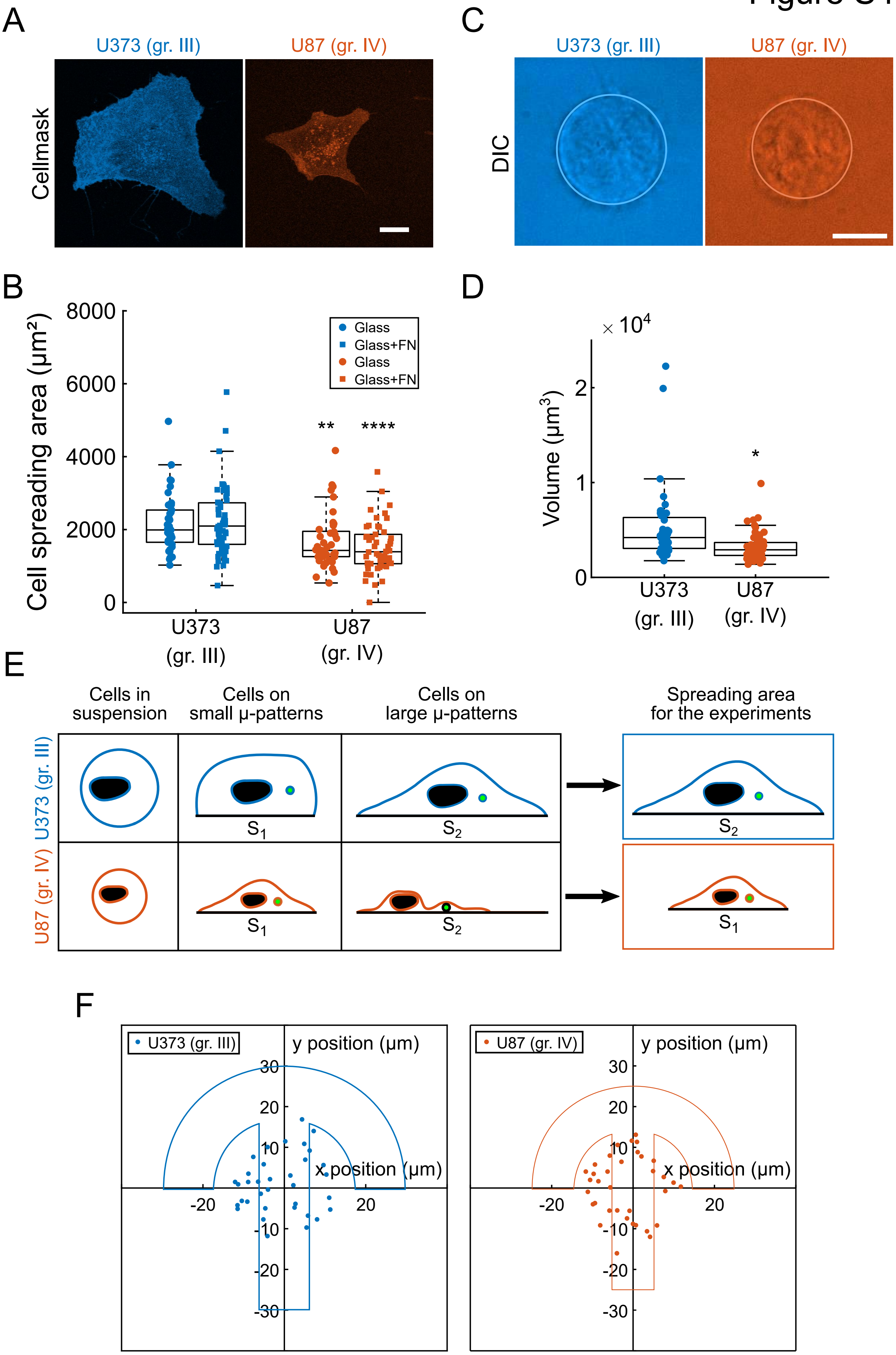

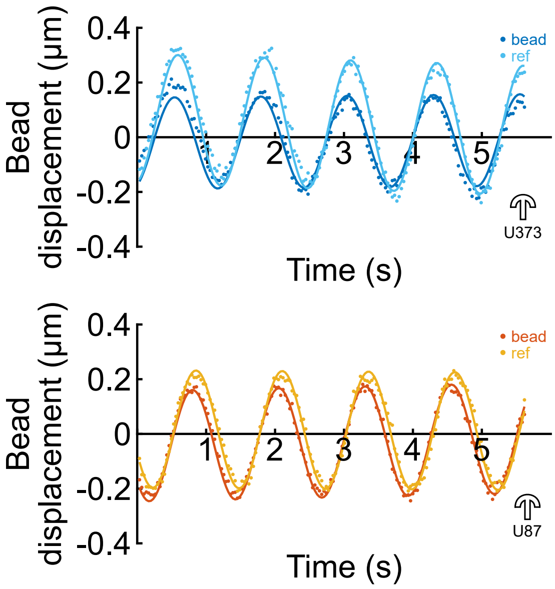

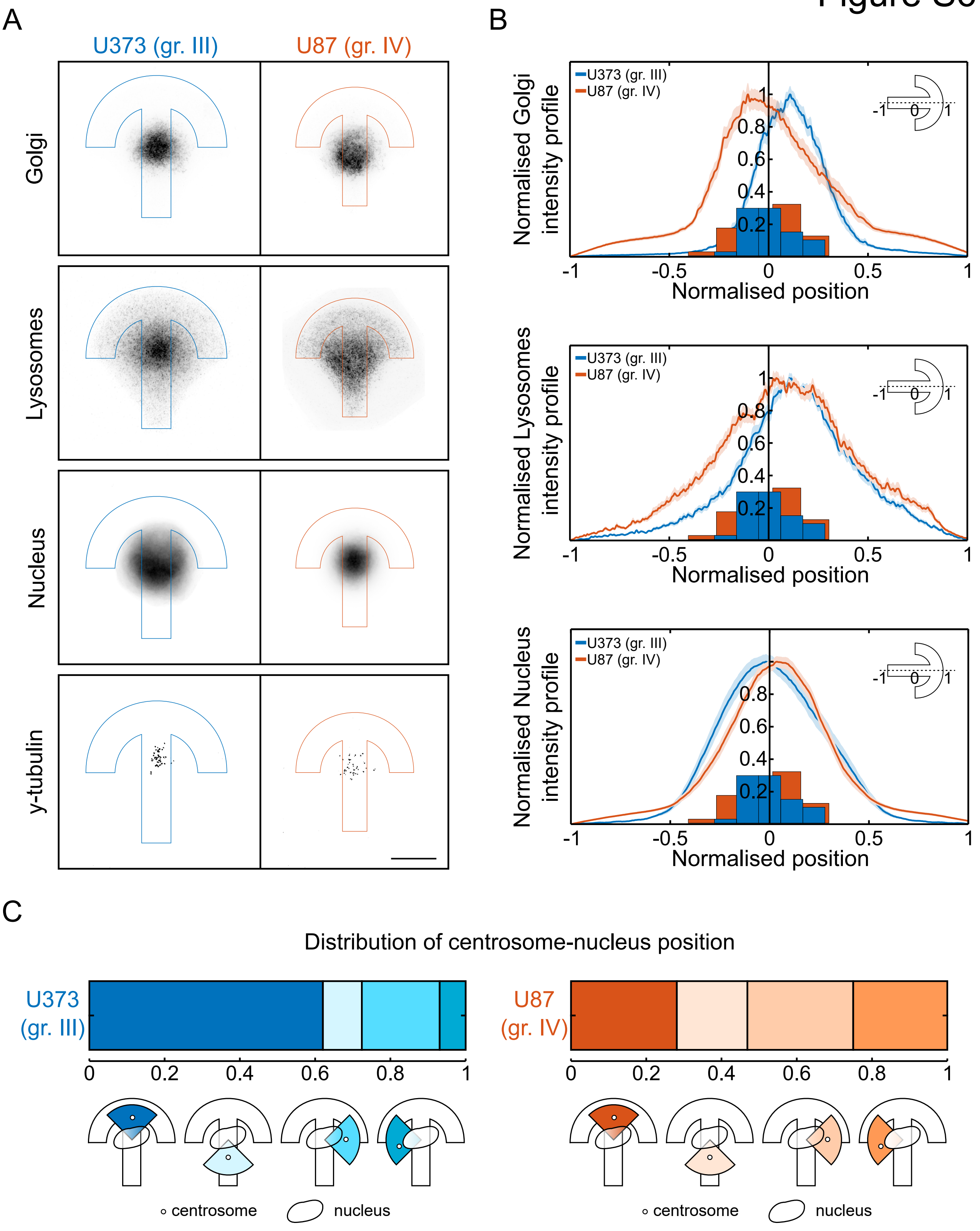

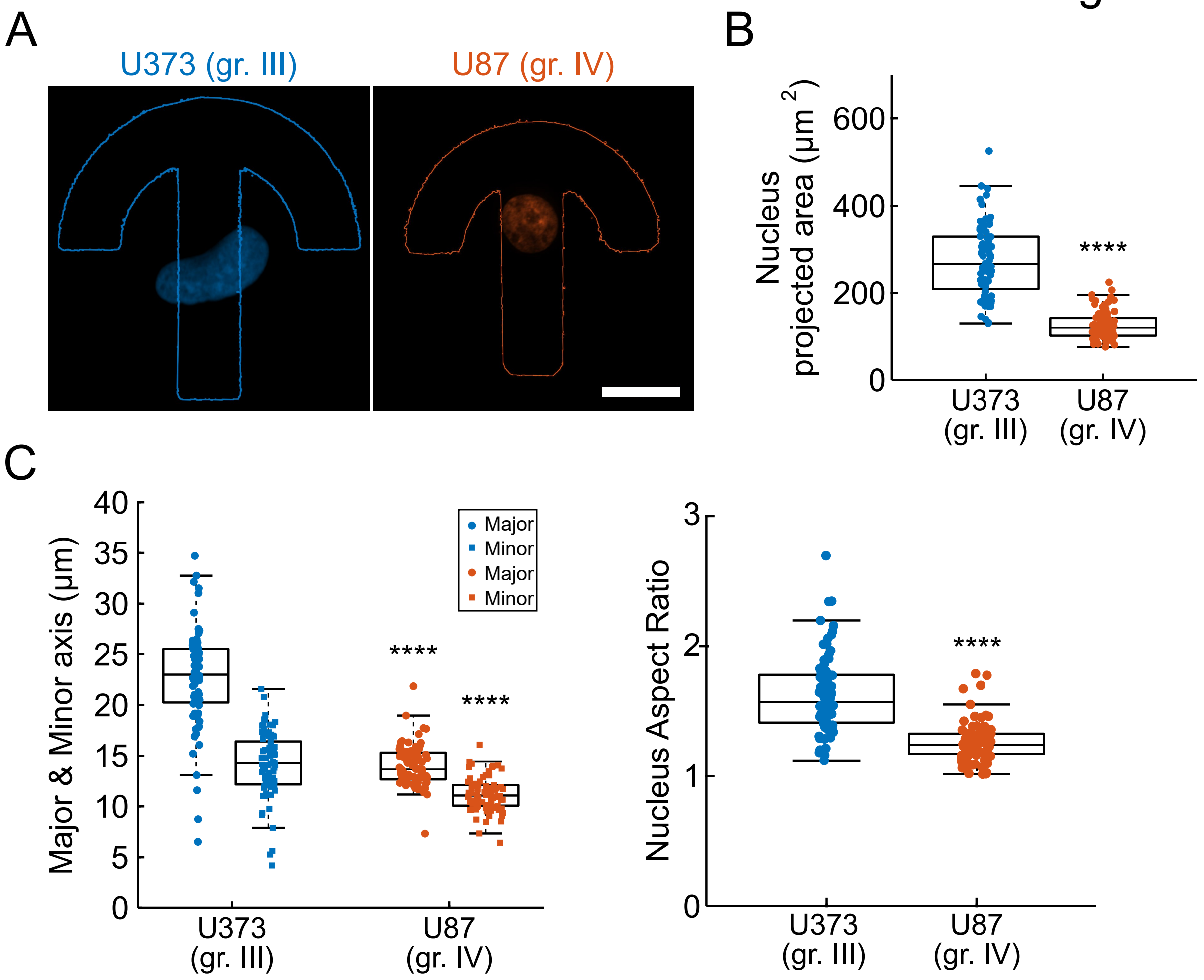
